# Pupil responses indicate selection of task-relevant background sounds during a dual continuous listening task

**DOI:** 10.1101/2025.05.20.655069

**Authors:** Lorenz Fiedler, Ingrid Johnsrude, Dorothea Wendt

## Abstract

Auditory attention can be voluntarily directed towards a sound source or automatically captured by background sounds, which may be either relevant, such that the listener shifts their attention to them, or irrelevant such that the listener tries to ignore or inhibit them. The ability to switch focus to a relevant sound source while inhibiting an irrelevant one requires attentional control and is crucial for navigating busy auditory scenes. Objective measures of attentional control could be beneficial in clinical contexts, such as fitting hearing aids. In a dual-task paradigm, we investigated whether pupil responses reflect relevance-dependent attentional selectivity. Participants with self-reported normal hearing (N = 21, Age: 27 to 66 years, pure tone average: −4 to +26 dB HL) listened to continuous speech from the front (primary task) while background sounds, consisting of cue names followed immediately by two-digit numbers, were presented from the left and right. The participant was told that one side, either right or left, was relevant and the other, irrelevant. The secondary task involved memorizing and later recognizing numbers from the relevant side. We observed increased pupil responses to sounds from the relevant side compared to the irrelevant side, indicating selectivity. Exploratory analysis showed that participants who exhibited stronger selectivity recognized more numbers correctly. Interestingly, pupil responses did not differ between hits and misses, but a stronger response to stream confusions versus correct rejections was found, suggesting that participants were more challenged by inhibiting irrelevant sounds than shifting attention to relevant sounds. In sum, our findings demonstrate that pupillometry provides valuable insights into attentional control abilities.

## Introduction

Humans use their sense of hearing not only to understand speech, but also to gain the situational awareness needed to navigate busy scenes. In both domains, attention plays a crucial role: On one hand, attention needs to be focused on features that separate target speech from noise (further called attentional allocation). On the other hand, occasionally attention must be switched to an alternative, relevant, sound source (further called attentional shifts), while irrelevant sounds should be ideally ignored (further called inhibition of attention). Balancing these three aspects of attentional navigation requires attentional control (Baddeley, 1996; Eysenck et al., 2007; Van Der Wel & Van Steenbergen, 2018), which has been studied extensively in the visual domain (for review: Goldstein et al., 2014). Attentional control abilities vary among individuals (Miyake et al., 2000). For example, some may have a harder time than others allocating sustained attention or inhibiting irrelevant information. This variability is an important consideration in clinical contexts (Wainstein et al., 2017), since different attentional deficits may require different treatment. In audiological contexts it may be relevant for hearing aid fittings as well, since a modern hearing aid is an adjustable directional and spectral filter, but existing fitting rules do not take attentional abilities into account.

An ideal hearing aid should restore all aspects of healthy hearing and support both comprehension of speech-in-noise and attentional control. Over the last few decades, the development of hearing aids has focused on optimizing speech-in-noise performance (Bentler, 2020). To this end, features for maximizing the signal-to-noise ratio have been developed. One of these features is directional filtering, which attenuates incoming sound signals from all directions except the front. Although the assumption of a frontal target is valid in many listening situations, it may not be valid when omnidirectional awareness is crucial, for example when navigating traffic. Indeed, besides facilitating the perception of target speech in background noise, hearing aid users require access to other priority sound sources in their surroundings (Byrne & Noble, 1998; Gatehouse & Noble, 2004; Nielsen & Henriksen, 2022). Hence, hearing aids should be optimized for situational awareness and tailored to individual attentional control abilities. However, standardized paradigms to assess auditory attentional control do not exist.

Sounds are thought to become the focus of attention in two ways, distinguished by the listener’s overarching intention (Kahneman, 1973): On one hand, voluntary attention (also called intentional or top-down attention) describes the act of actively focusing attention on a target. For example, sustained allocation of attention to an audiobook can be seen as an active decision and thus falls under this category. On the other hand, stimulus-driven attention (also called involuntary, bottom-up or automatic attention) refers to the fact that attention can be captured by a stimulus outside the current focus, for example by someone calling our name (Cherry, 1953). Although the initial capture of attention may be automatic, the relevance of the attention-capturing stimulus determines whether voluntary attention follows, resulting in an intentional attentional shift, or whether distraction is kept to a minimum (i.e., the transiently distracting stimulus is subsequently inhibited). Both of these subsequent reactions, based on the relevance of the interrupting sound to the listener’s current goal, require attentional control.

Since attention is closely linked to cognitive effort (Bruya & Tang, 2018; Hess & Polt, 1964; Kahneman, 1973) and pupillometry has been found to indicate cognitive effort (Kahneman & Beatty, 1966), the amount of deployed attentional resources can be studied using pupillometry (Hoeks & Levelt, 1993; Iriki et al., 1996; Smallwood et al., 2011; Van Der Wel & Van Steenbergen, 2018; Wierda et al., 2012). In the auditory domain, pupillometry has been extensively used to study listening effort (Kadem et al., 2020; Kramer et al., 1997; Ohlenforst et al., 2017, 2018; M. B. Winn et al., 2015; M. B. Winn, 2016). While listening effort can be seen as the total effort exerted for executive functions required for listening (prominently to speech), some of these executive functions are certainly orchestrated by attention. Accordingly, the pupil response has been found to be stronger to task-relevant sentences compared to task-irrelevant ones (Pielage et al., 2021). The pupil response to irrelevant background sound indicates the degree of distraction (Fiedler et al., 2025). In sum, pupil responses may be well suited to monitor the selectivity in the deployment of attentional resources in a continuous listening task, in which attentional allocation, attention shifting and inhibition of attention are all required.

Cognitive effort can lead to both phasic and tonic changes in pupil size (Beatty, 1982). The prototypical pupil response to a discrete and cognitively demanding stimulus is a dilation with a peak after about 1.5 seconds and a subsequent return back to baseline about 3 seconds (Hoeks & Levelt, 1993). Such a response can be considered a phasic response, because it has no longer lasting effects on the pupil size. It indicates how much attentional resources are allocated to the processing of a stimulus, both stimulus-driven and voluntarily. Hence, a phasic response can be expected both in response to irrelevant and relevant stimuli, while both higher degrees of stimulus-driven attention and task-relevance should lead to greater pupil dilation, respectively. On the other hand, tonic changes are longer lasting offsets in pupil size, for example caused by anticipatory arousal (Cole et al., 2022) or memory load (Bönitz et al., 2021; Kahneman & Beatty, 1966). Memory-related tonic changes become relevant when a sustained task requires memory maintenance, which we would only expect in the case of relevant stimuli. Phasic and tonic pupil responses have in common that a stronger dilation indicates higher cognitive effort.

We designed an auditory attentional control task where participants were asked to allocate sustained attention to an audiobook (primary task), while occasionally shifting attention towards relevant background sounds (secondary task), and ignoring irrelevant ones. We hypothesized a relevance-dependent response in the pupil size: Relevant background sounds should lead to greater pupil dilation. Since behavioral results alone would not indicate whether errors in secondary task performance are driven by unsuccessful shifting or unsuccessful inhibition, in an exploratory analysis we contrasted the pupil responses both to relevant sounds (*hits* vs. *misses*) and to irrelevant sounds (*stream confusions* vs. *correct rejections*). Intentionally selected background sounds (*hits* and *stream confusions*) should lead to greater pupil dilation than their non-selected counterparts (*misses* and *correct rejections*).

## Materials & Methods

### Participants

Twenty-one participants with self-reported normal hearing (9 female, 12 male) were invited to a single visit lasting approximately 2.5 hours length. Their age ranged from 27 to 66 years (mean: 47.3; SD: 13.3). Hearing thresholds were measured and the pure-tone average (PTA) between 500 and 4000 Hz ranged from −3.93 to 26.1 dB HL (mean: 9.12; SD: 7.44). This indicates that some participants’ hearing thresholds were in the range of clinically mild hearing loss, but the majority had clinically normal hearing. As expected, age and PTA positively correlated (Fig. 1A, r = 0.570, p = 5.90×10^-3^).

**Figure 1:**
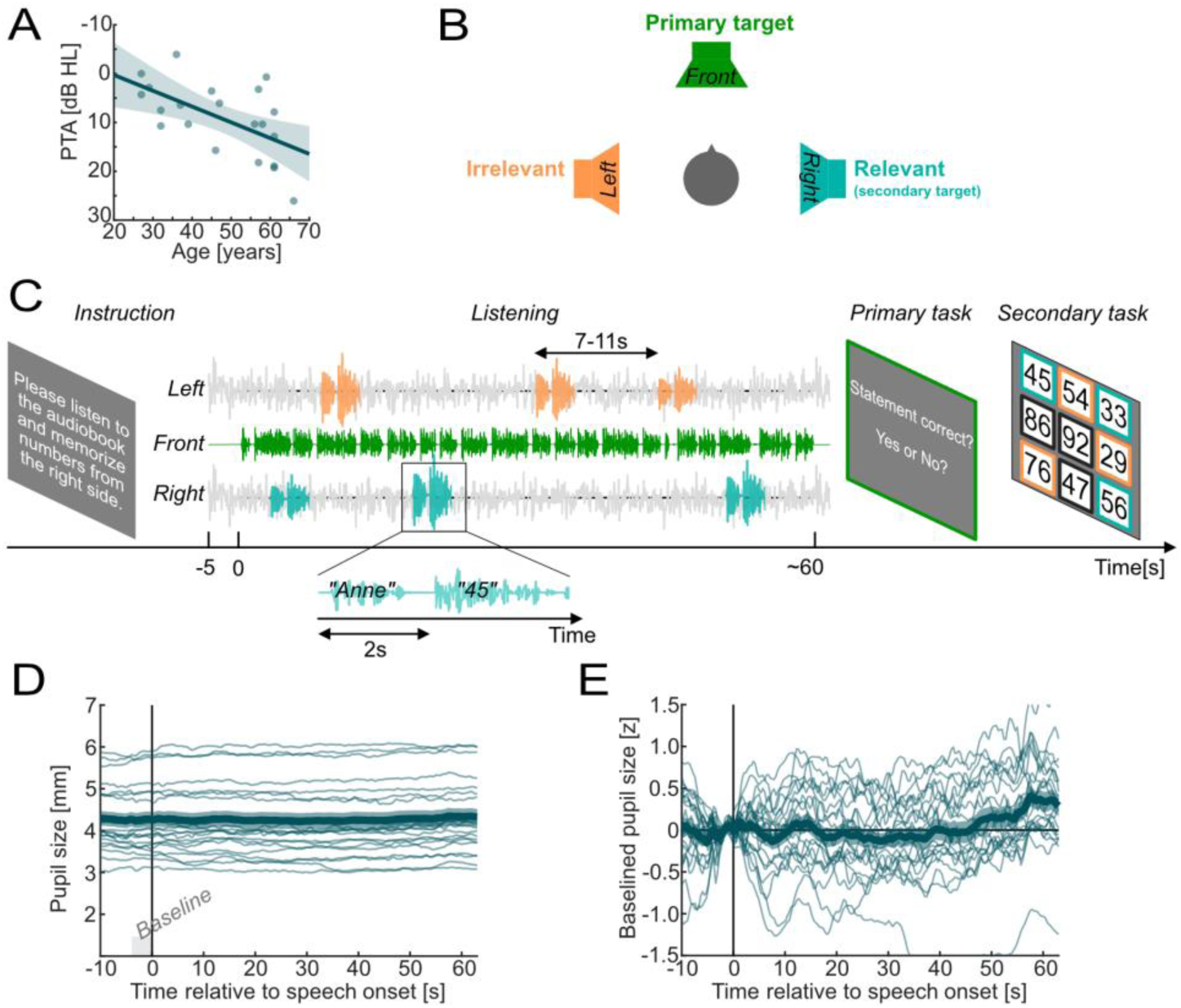
Participants and task design. **A)** Scatter plot of age and pure-tone average (PTA) with fitted least-squares line and shaded area depicting the 95% confidence band. **B)** Spatial setup: Three loudspeakers (front: 0, right: −90, left: +90 degrees). The primary target speech (audiobook; green) was presented from the front. Task-relevant secondary targets (names & numbers; cyan) were presented from left or right (balanced across trials), while task-irrelevant distractors (names & numbers; orange) were presented at the opposite side. **C)** Example trial: Visual instruction to listen to the audiobook and memorize numbers from the right (or left) side. After starting the trial with a button press, 5 seconds of silence were followed by 5 seconds of babble noise, followed by the primary target speech of around 1 minute. Relevant (cyan) and irrelevant (orange) background sounds consisting of a name followed by a number after 2 seconds were presented in random order and at random points in time. After two comprehension questions participants were asked to pick the 3 relevant numbers from a set of 9. Participants had to pick exactly 3 numbers, which could result in hits (i.e., cyan items), stream confusions (i.e., orange items) or random errors (i.e., black items). **D)** Pupil size across trial. **E)** Pupil size across trial z-scored based on baseline (−3 and 0 sec). For D & E: Thick lines and shaded area depict mean across participants ±SEM. Thin lines depict individual means across trials.

The sample size was based on a power analysis with G*power (Faul et al., 2007) on data from a pilot study with 7 participants. Our main effect of interest (the selectivity in the pupil response) reached an effect size of Cohens d = 1.27 which under a power of 1 - β = 0.95 and a significance level of α = 0.05 results in a recommended sample size of 11. We were also interested in the behavior-dependent modulation of the main effect, but we could not get a proper estimate of its effect size due to insufficient number of behavioral errors in the pilot data. We decided to increase the power and aimed at a sample size of around 20 participants. However, since the effect size (and hence the statistical power) was unknown, we consider the additional analysis exploratory.

The experiments were conducted at Eriksholm Research Centre (Snekkersten, Denmark). The study was approved by the Copenhagen regional ethics committee (De Videnskabsetiske Komitéer, Region Hovedstaden).

### Apparatus

The hardware setup was identical to the one used in Fiedler et al. (2025). The experiment took place in an acoustically and electro-magnetically shielded booth. A comfortable office chair was placed at a reference point in the middle of the room in front of a desk. A computer keyboard and a mouse were placed on the desk. The left and right Ctrl-buttons of the keyboard were labelled with “Ja” & “Nej” (Yes & No). The assignment of the labels to left and right was balanced across participants. The space bar was labelled with “Start”. Behind the keyboard, a Tobii Pro Spectrum eye tracker (Tobii AB, Stockholm, Sweden) was placed. The light conditions were kept at a moderate level and measured at 75 lux at the reference point.

Three loudspeakers (Genelec 8030A, Genelec Oy, Iisalmi, Finland) were placed one meter from the reference point. One loudspeaker was positioned at 0 degrees azimuth (front, behind the desk, above the screen), and two were positioned to the left and right at ±90 degrees azimuth. The loudspeakers were connected to an RME Multiface II audio interface (AudioAG, Haimhausen, Germany). The audio interface was connected via USB to a personal computer running Microsoft Windows 10 (Microsoft Corp., Redmond, USA), where MATLAB 2020b (Mathworks Inc., Natick, USA) and Tobii Eye Tracker Manager were installed. The audio was controlled from MATLAB using Soundmexpro toolbox (Hörtech gGmbH, Oldenburg, Germany). Underneath the frontal loudspeaker, a screen was placed to display task instructions and comprehension questions. The screen was controlled via MATLAB using Psychtoolbox version 3.0.17 (Brainard, 1997). A BioSemi ActiveTwo EEG amplifier (BioSemi B.V., Amsterdam, The Netherlands) was placed behind the participant. Further details on EEG data will be reported elsewhere.

### Stimulus material

The goal was to present continuous speech stimuli as the primary target along with relatively short stimuli used as both relevant (i.e., secondary target) and irrelevant (i.e., distractor) background sounds embedded in unintelligible noise.

An audiobook was prepared as the primary, continuous speech target. We used a biography about a controversial historic Danish person called Simon Spies (John Lindskog, “Simon”) spoken by a male voice, as used in an earlier study (Borges et al., 2025). The audiobook was cut into 46 parts (6 for training + 40 for the main experiment), each of at least one minute in length, while making sure that narrative phrases stayed intact, resulting in a variable length between 60.0 and 72.3 seconds (mean: 63.8; SD: 2.56).

Spoken names and numbers were prepared as background sounds. While names served as a cue, numbers were used as secondary targets and distractors (see Procedure & task, below). Both names and numbers were generated using online text-to-speech software (www.Narakeet.com) with the Danish preset voice “Trine”. The 126 most common names in Denmark in the year 2022 (63 male, 63 female) were used (these were provided by Danmarks Statistik (www.dst.dk) upon request). Identically pronounced names (e.g., Caroline & Karoline) were treated as identical. Two-digit numbers between 21 and 99 excluding 30,40,50 etc. were used.

Babble noise was created within which the secondary target and distractor items would be embedded. We used Danish radio news clips, which had been earlier used as target stimuli in previous research (Alickovic et al., 2020, 2021; Fiedler et al., 2021). The collection consisted of a total of 320 newsclips spoken by four different talkers (2 female, 2 male, 80 newsclips each). A total of 44 babble noise files were created (4 for training + 40 for the main experiment, 81 seconds each) by randomly concatenating and mixing newsclips into two channels with the criterion that a mix of the 4 different talkers occurs in each channel and that the same newsclip did not occur in both channels at the same time. As a result, we obtained 8-talker babble noise distributed across two channels.

All stimuli were normalized to an equal root-mean-square (RMS). A hearing threshold was determined for each listener for the audiobook by presenting sections of the audiobook at an increasing level (2 dB per second) and asking the participants to press a button as soon as they could hear the slightest whisper. After eight repetitions, the average level during button press was used as the individual, stimulus-specific hearing threshold.

The audiobook was calibrated to 50 dB above the individual hearing threshold, resulting in an average presentation level of 62.8 dBA (SD: 6.5 dB). The RMS-equalized name and number stimuli were calibrated to −3 dB relative to the audiobook. Each of the two babble noise channels was calibrated to −15 dB relative to the audiobook, so the summed level of the two channels was −12 dB relative to the audiobook.

### Procedure & task

Before the visit, participants were asked to read a document about the procedure. At the beginning of their visit, participants gave written informed consent. The whole visit took approximately 3 hours. First, a pure-tone audiogram was conducted. Next, the participants were fitted with EEG sensors for another study (results not reported here). The eye tracker was adjusted to reliably capture both eyes and calibrated using the five-dot calibration procedure in Tobii Eye Tracker Manager. The individual, stimulus-specific hearing threshold was determined as described above (see Stimulus material).

An example trial of the main experiment is depicted in Figure 1C. First, an instruction appeared on the screen asking the participant to listen to the audiobook (i.e., primary task) and memorize numbers from the left or right side (i.e., secondary task). The assignment of the relevant side was balanced across the experiment and randomized trial by trial. Participants started each trial by pressing the space bar. The screen turned black, and participants were asked to look at the loudspeaker in front of them. Babble noise began at the left and right loudspeaker after 5 seconds of silence. For each participant, babble noise files (see Stimulus material) were randomly permuted across trials. Another 5 seconds later the audiobook began, which lasted for at least 60 and up to 72.3 seconds (see Stimulus material). The audiobook was presented in its original order to allow participants to follow the plot.

In each trial while the audiobook was presented, 6 name-number pairs (i.e., background sounds) were presented (3 on each side). There was a two-second gap between the onset of the name and the subsequent number. The name-to-name interval was jittered by permuting a uniform distribution between 7 and 11 seconds in steps of 0.8 among the 6 background sounds. The assignment of names to the relevant and irrelevant side was balanced across the experiment, ensuring each name occurred once on the relevant and once on the irrelevant side. Within a trial, a particular name or number did not occur more than once, and the order of relevant and irrelevant name-number pairs was randomized. Babble noise continued for 3 seconds beyond the audiobook.

After the babble noise stopped, participants were prompted with two statements about the content of the audiobook. They were asked to indicate whether a statement was true using the computer keyboard buttons labelled with “Ja” (Yes) or “Nej” (No). Subsequently, participants had to pick exactly 3 numbers from a 3-by-3 grid of nine numbers. They were asked to select the 3 relevant numbers if they could recognize them or to make a guess otherwise. Besides the 3 relevant numbers, the 3 numbers from the irrelevant side were offered as well as 3 numbers that did not occur during the trial. Hence, the behavioral outcome in the secondary task could either be a *hit* if a relevant number was selected, *stream confusion* if an irrelevant number was selected, or *random error* if a non-presented number was selected.

The experiment started with six training trials to introduce participants to the task in a stepwise fashion: In the first two trials participants were introduced to the primary task. In training trials 3 & 4, babble noise was added. In trials 5 & 6, participants were introduced to the secondary task. In the main experiment participants underwent 40 trials split up into four blocks of 10 trials each (approximately 15 minutes). Between each block, participants were offered a break.

### Pupillometry

The preprocessing of pupillometric data was done identically to Fiedler et al. (2025). We recorded pupillometric data at a sampling rate of 1200 Hz throughout the trial. The raw pupillometric data, measured in millimeters, contained NaN-values at time points where the pupil could not be detected (i.e., missing values) and was cleaned of artifacts (M. B. Winn et al., 2018). When these missing values exceeded a duration of 20 milliseconds (i.e., 24 samples) we extended the duration of missing values to include 40 samples (35 milliseconds) prior to the onset of the original NaN string, and 120 samples (100 milliseconds) after its offset. This longer duration of missing values increased our chance of eliminating artefacts caused by the failure of pupil tracking, and by resumption of tracking. Remaining spikes in the data were removed based on the criterion that the sample-to-sample difference must not exceed two standard deviations as calculated from the same trial. If this criterion was met, both samples were replaced with NaN-values. All missing values were linearly interpolated and the pupillometric data were low-pass filtered by a convolution with a one-second-wide Hamming window. As an intermediate step, we obtained one continuous pupil trace for each trial (Fig 1D).

Previous studies found that age and PTA can influence both baseline and the dynamic range of the pupil size (Fiedler et al., 2025; B. Winn et al., 1994). To rule out any related confounds, the individual pupillary data were z-scored based on parameters (mean and standard deviation) extracted from the baseline interval between −3 and 0 seconds relative to speech onset. To this end, all baseline periods were concatenated, and the two parameters were estimated once for each participant. The pupillary data of each participant were then z-scored by subtracting the individual mean and dividing by the individual standard deviation (Fig. 1E). Note however, z-scoring did not have a substantial impact on the results and similar results would have been obtained even without z-scoring.

To extract the pupil size time course around the time of presentation of the background sounds, the preprocessed pupil size data were epoched between −1 and 8 seconds relative to each name onset. The mean of the pupil size in the baseline interval between −1 and 0 seconds was subtracted from the pupil size time course. We kept only pupil responses in which less than 30% of the values were missing before interpolation. For each participant, the eye with more available responses was picked for further analysis. We obtained pupil responses to an average of 87.4% of the background sounds. More than 80% of the pupil responses were available for 18 participants, while for the other three participants only 49.2%, 39.2%, and 37.9% were available. All available data were used for the subsequent analysis. There was no significant difference between the number of pupil responses available to relevant vs. irrelevant background sounds (two-sided paired t-test, t(20) = 0.0553, p = 0.956).

### Statistical analysis

#### General statistical approach

In case of within-subject comparisons, we applied repeated-measures parametric statistics: In case of paired comparisons two-sided, paired-samples t-tests were used and effect sizes will be reported as Cohens d. In case of multi-factorial comparisons, repeated-measures ANOVAs were used and effect sizes will be reported as partial η^2^.

In case of time series analysis a cluster-based permutation test with 5000 permutations was performed (Maris & Oostenveld, 2007). In brief, a cluster-test detects time windows of significant differences based on an underlying parametric test at each time point, while controlling for multiple comparisons. First, clusters of contiguous timepoints with significant test statistics were detected in the original data. The test statistics were summed across each cluster. Second, the condition labels of the data were randomly permuted 5000 times and within each permutation, clusters of contiguous timepoints with significant test statistics were detected and the test statistics were summed across each cluster. The reported probability p_perm_ refers to the rate of random permutations in which at least one cluster with an equal or higher test-statistic sum has been found compared to each cluster in the original data, respectively. Subsequently, a single parametric test was done on the means across the detected original clusters in order to obtain effect size measures (Cohens d or partial η^2^).

In case of between-subject comparisons Pearson correlation was used. Note that correlation analyses were either used to detect confounds within the sample or to provide exploratory analysis. Since the sample size does not allow a precise estimate of the true correlation coefficient of the population (Schönbrodt & Perugini, 2013), 95%-confidence bands (CI_r_) will be reported alongside with the correlation coefficients of the sample.

#### Behavioral and demographic data

The behavioral performance in both the primary and secondary tasks was analyzed. In the primary task the answer could be either correct or wrong with a chance level of 50% and the individual performance will be referred to as *correct answer rate*. The secondary task performance was divided into *hits*, *stream confusions*, and *random errors* with chance level of 33.3%. To estimate whether irrelevant background sounds were indeed distracting (Wöstmann et al., 2022), a t-test was performed under the hypothesis that participants made more *stream confusions* than *random errors*.

To extract a single behavioral measure for the secondary task that captures the selectivity between relevant and irrelevant background sounds, the difference between the rates of *hits* and *stream confusions* was calculated and will be further called *behavioral selectivity*. This measure takes into account that a certain performance level in the secondary task depends on both the ability to switch attention to relevant background sounds and inhibit irrelevant ones. While two individuals may show the same amount of *hits*, one may show a higher number of *stream confusions* than the other, which would result in lower *behavioral selectivity* for the first participant compared to the second. To estimate whether participants prioritized the tasks differently, the correlation between *correct answer rate* and *behavioral selectivity* was calculated.

A control analysis was performed to determine whether *stream confusions* were caused by mistakenly considering the irrelevant side as the relevant one. If this were the case, stream confusion errors should cluster within trials. Clustering would manifest as a significant deviation from a random distribution of stream confusions across trials. To test this, surrogate data were created by randomly shuffling the secondary task behavioral data 1000 times within each participant and counting the number of stream confusions per trial. Within each of the 4 bins of 0 to 3 possible stream confusions, a t-test was used to test systematic deviations from the random distribution. If clustering occurred, we would expect to see an increased count in the bins for 2 & 3 stream confusion and a reduced count in the bins for 0 and 1 stream confusions. In a descriptive analysis, age and PTA were Pearson-correlated with *correct answer rate* and *behavioral selectivity*, respectively.

#### Pupillary data

We refer to contrasts between pupil responses to relevant and irrelevant background sounds as *pupil selectivity.* First, to estimate the time range of *pupil selectivity,* a cluster-based permutation test was performed (see details in General Statistical Approach, above) in the time range between 0 and 5 seconds relative to name onset with an underlying t-test. The retained cluster (see results) was used to calculate the mean pupil dilation (MPD). To confirm *pupil selectivity*, MPD for relevant and irrelevant sounds was contrasted using a t-test.

In an exploratory analysis, we estimated whether the behavioral fate of background sounds could be predicted by pupil responses. To this end, pupil responses were categorized into four groups: A relevant background sound could either end up as a *hit* or a *miss*, while an irrelevant background sound could either end up as a *correct rejection* or a *stream confusion*. First, for relevant background sounds, we expected a smaller pupil dilation for a *miss* compared to a *hit*. Second, for irrelevant background sounds, we expected smaller pupil dilation for a *correct rejection* compared to a *stream confusion*. In other words, we expected greater pupil dilation in response to background sounds that were selected (*hits* and *stream confusions*) than to background sounds that were not selected (*misses* and *correct rejections*). Accordingly, we ran a cluster-based permutation test in the time range between 0 and 5 seconds relative to name onset with an underlying 2×2 ANOVA. The first factor was *Relevance* with the levels *relevant* and *irrelevant*. The second factor was *Selection* with the levels *selected* (*hits* and *stream confusions*) and *non-selected* (*misses* and *correct rejections*). Note that one participant performed at 100% in the secondary task and thus did neither produce *misses* nor *stream confusions,* so they were excluded from this part of the analysis.

In another exploratory analysis to investigate whether strong *pupil selectivity* correlates with better secondary task performance, the individual *pupil selectivity* was Pearson-correlated with *behavioral selectivity*. To confirm that the *pupil selectivity* was not only driven by the number of stream confusions (see results for more details), we repeated the Pearson-correlation between the *pupil selectivity* and *behavioral selectivity*, but this time only for responses that ended up as *hits* or *correct rejections*.

In a final control analysis we examined how constant both behavioral performance and pupil responses were across the four blocks of the experiment, and across a trial. To this end, we binned behavioral performance (both primary and secondary) and MPD within each of the four successive blocks, and secondary task performance and MPD into background sound serial position (1-6) within trials. With individual linear least-square fits and one-sample t-tests of the slope estimates against zero, we estimated whether both behavioral performance and pupil responses were constant or whether there was a significant deflection from zero.

## Results

### Behavioral performance

On the primary task, all participants performed well above chance (Correct Answers; Fig. 2A). Participants demonstrated strong comprehension, answering 80.6% of the questions correctly (SD: 4.04%, range: 72.5 – 87.5%).

**Figure 2:**
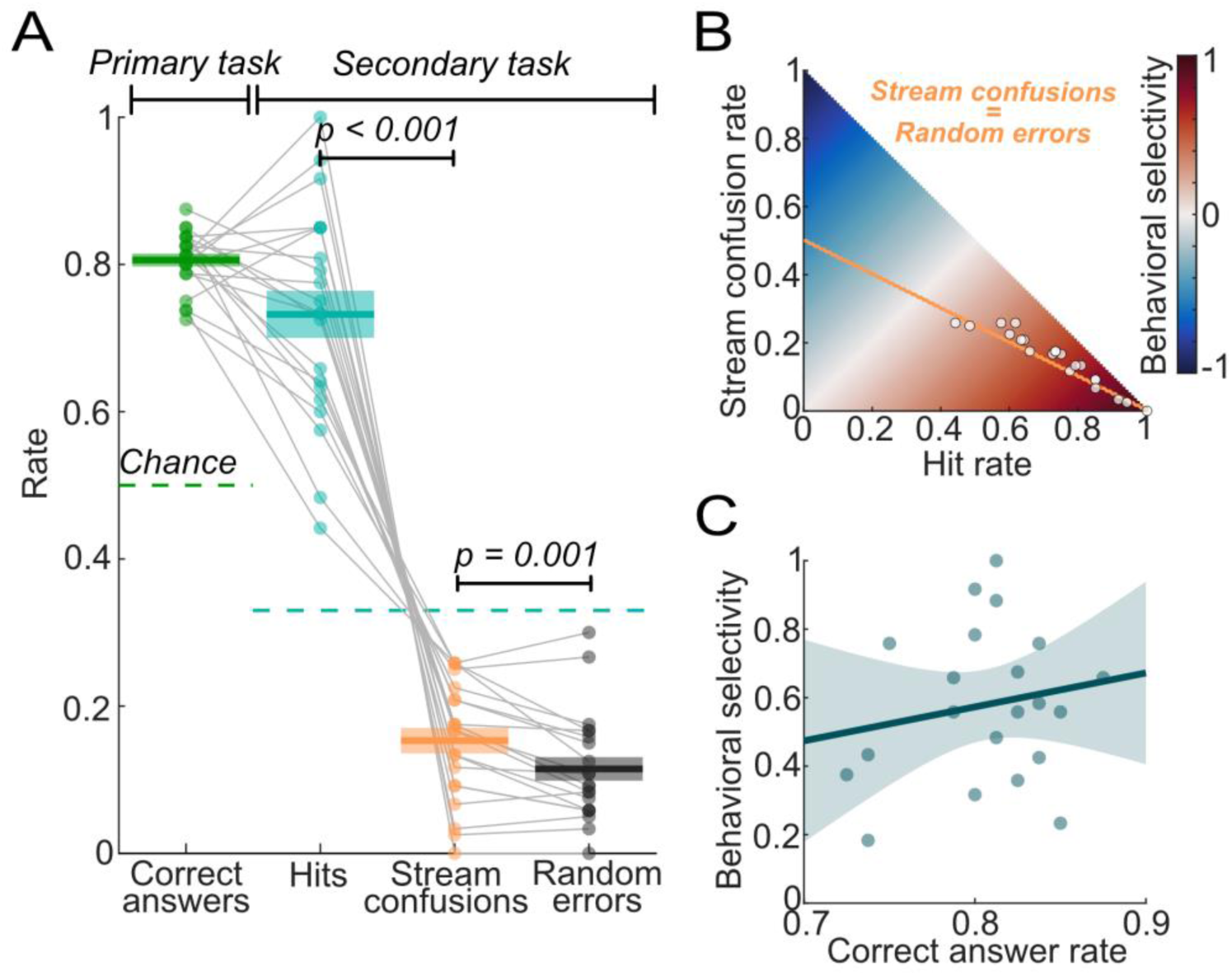
Behavioral performance. **A)** Primary task performance as proportion of correctly answered questions and secondary task performance as proportion of hits as well as errors divided into stream confusions and random errors. Thick solid lines embedded in shaded areas depict the mean ±1 standard error. Dots depict individual means. Thin grey lines connect data points of the same subject. **B)** Behavioral selectivity was calculated as the difference between hits and stream confusions. Orange line marks equal amount of stream confusions and random errors. White dots depict individual rates. **C)** Scatter plot of primary task performance (correct answer rate) and secondary task performance (behavioral selectivity) with fitted least-squares line and shaded area depicting the 95% confidence band.

On the secondary task, hit rates were also above chance, but were more variable compared to the primary task (Hits; Fig. 2A). On average, participants correctly picked 73.2% of the numbers from the relevant side (SD: 14.6%, range: 44.2 – 100%). Errors were more likely to involve selecting numbers from the irrelevant side (Stream confusions; Fig. 2A) rather than numbers not presented during the trial (Random errors; Fig. 2A), which was confirmed by a two-sided paired t-test (t(20) = 3.86, p = 9.72×10^-4^, Cohens d = 0.843) and is also illustrated in Figure 2B. On average, participants made 15.3% stream confusions (SD: 7.97%, range: 0 – 25.8%) and 11.47% random errors (SD: 7.35%, range: 0 – 30%). To obtain a behavioral performance measure that takes both hits and stream confusions into account, *behavioral selectivity* was calculated as the difference between *hit rate* and *stream confusion rate* (Fig. 2B). There was no significant correlation between *correct answer rate* and *behavioral selectivity* (r = 0.179, CI_r_ = [-0.274; 0.567], p = 0.437, Fig. 2C).

In sum, the behavioral performance suggests that all participants were able to adequately perform the dual task at above chance levels, but the secondary task challenged them to varying degrees. This variability could neither be explained by age, PTA (see control analysis below), nor primary task performance.

### Selective pupil response between relevant and irrelevant sounds

We analyzed the pupil response across time relative to the onset of names followed by a number two seconds later. Pupil responses were contrasted based on the secondary task instruction, which rendered names and numbers either relevant or irrelevant. We call the difference in pupil size for relevant, compared to irrelevant, background sounds “pupil selectivity”.

Figure 3A depicts baselined pupil size relative to name onset. There was a clear, step-like increase in pupil size in response to relevant background sounds, while the response to irrelevant counterparts was less pronounced. Interestingly, the pupil size in response to relevant sounds did not return to baseline during the period of interest, while the pupil size following irrelevant sounds showed a negative drift below baseline.

**Figure 3:**
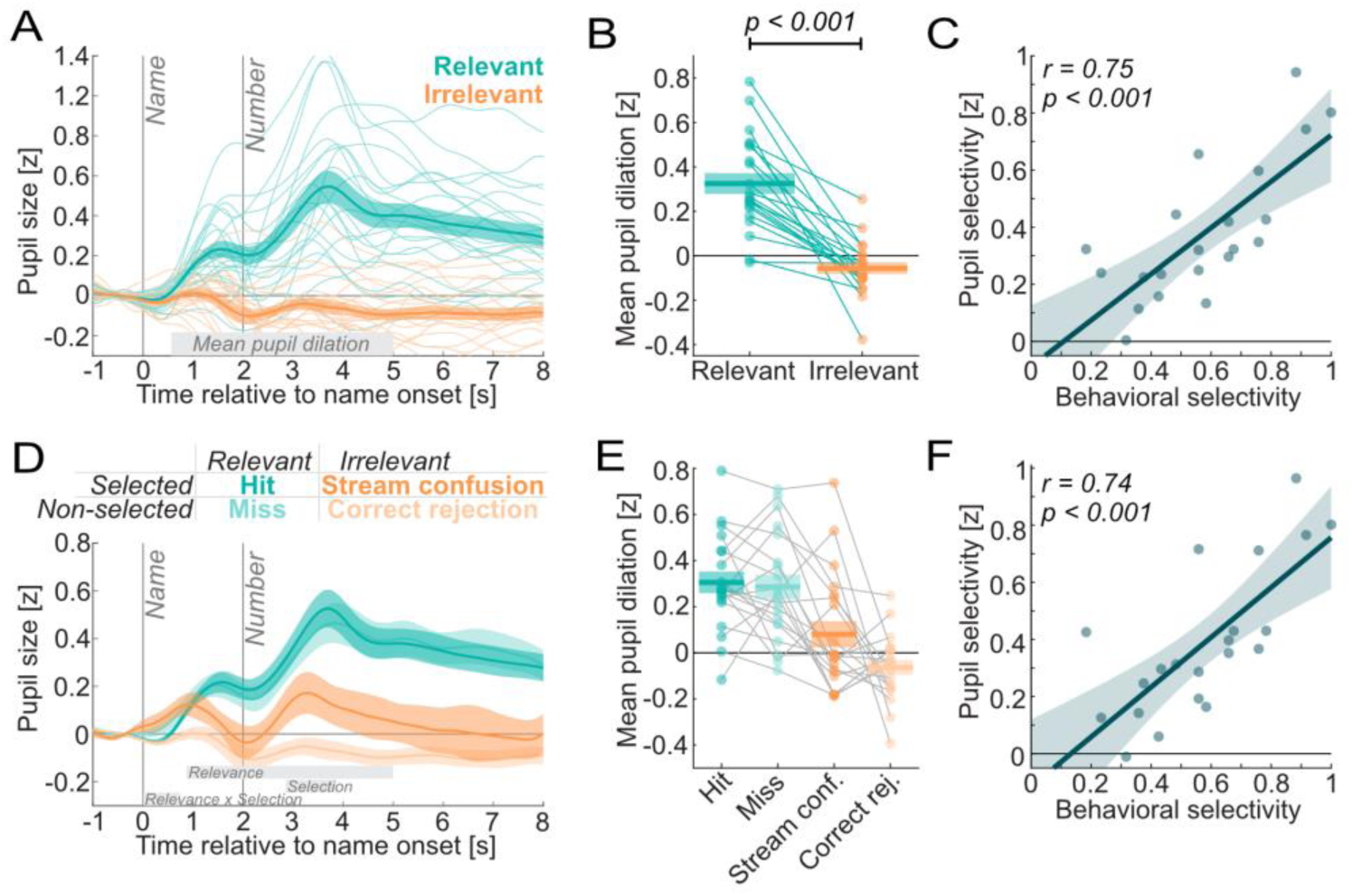
Pupil response to background sounds. Thick solid lines embedded in shaded areas depict the mean ±1 standard error. Dots and thin lines depict individual means. Thin grey lines connect data points of the same subject. **A)** Pupil responses to relevant and irrelevant background sounds. Vertical grey lines indicate the onset of name and numbers. Grey bar indicates time window for mean pupil dilation. **B)** Mean pupil dilation (MPD; 0.53 to 5 seconds relative to name onset). **C)** Scatter plot depicts the relationship between the difference in MPD (i.e., pupil selectivity) and secondary task performance (i.e., behavioral selectivity) with fitted least-squares line and shaded area depicting the 95% confidence band. **D)** Pupil responses by behavioral outcome. Grey bars indicate significant clusters of main effects and interaction. **E)** MPD by behavioral outcome. **F)** Same as C, but only for responses that resulted in hits or correct rejections.

We analyzed *pupil selectivity* by contrasting pupil size between relevant and irrelevant background sounds between 0 and 5 seconds via a cluster-based permutation test. This revealed a cluster between 0.53 and 5 seconds (p_perm_ < 0.001). Accordingly, the mean pupil dilation across that time period showed a significant difference (mean = 0.378, SD = 0.241, t(20) = 7.18, p = 5.93×10^-7^, Cohens d = 1.567). Consistently, all participants’ pupils responded more strongly to relevant background sounds compared to irrelevant ones (Fig. 3B), but the strength of this selectivity varied among subjects. Next, we analyzed whether this *pupil selectivity* correlated with behavioral performance.

### Exploratory analysis: Selectivity in pupil response predicts secondary task performance

To find out to which extent the relevance-dependent pupil responses are linked to behavioral performance, the Pearson-correlation between *pupil selectivity* and *behavioral selectivity* was calculated (Fig. 3C). Participants with stronger *pupil selectivity* also showed better behavioral performance in the secondary task: *pupil selectivity* positively correlated with *behavioral selectivity* (r = 0.747, CI_r_ = [0.465; 0.891], p = 1.01×10^-4^).

### Exploratory analysis: Increased pupil response to stream-confused irrelevant sounds

Since we found a significant correlation between *pupil selectivity* and *behavioral selectivity*, we investigated whether *pupil selectivity* was driven by differences in pupil responses based on the behavioral fate of the background sounds.

Figure 3D depicts the pupil responses categorized into their behavioral outcome of *hits*, *misses*, *stream confusions,* and *correct rejections*. While there was only a slightly stronger response to *hits* compared to *misses*, a stronger response to *stream confusions* compared to *correct rejections* was observed. A cluster-based permutation test based on a repeated-measures ANOVA revealed several clusters: First, there was a cluster showing the main effect of *Relevance* between 0.87 and 5 seconds (p_perm_ < 0.001), which corresponds to the *pupil selectivity* cluster found earlier. Second, there was a cluster of a main effect of *Selection* between 2.87 and 3.95 seconds, which just failed to reach statistical significance (p_perm_ = 0.0728). However, a repeated-measures ANOVA on the means across the respective time period revealed a main effect of selection (F(1,19) = 5.70, p = 0.0275, partial η^2^ = 0.231). A t-test across the time period between the mean of *selected* and *non-selected* responses was significant as well (mean = 0.134, SD = 0.251, t(19) = 2.33, p = 0.0275, Cohens d = 0.534). Third, there was a cluster showing an interaction between *Relevance* and *Selection* between 0.025 and 0.84 seconds, which just failed to reach statistical significance (p_perm_ = 0.0832). However, a repeated-measures ANOVA on the means across the respective time period revealed a significant interaction (F(1,19) = 6.66, p = 0.0183, partial η^2^ = 0.259). A t-test across the time period between the relevance-related difference between *selected* and *non-selected* responses was significant as well (mean = 0.143, SD = 0.249, t(19) = 2.58, p = 0.0183, Cohens d = 0.577). In sum, there was a tendency towards an increased response to selected background sounds mainly driven by the stronger response to irrelevant sounds that were stream-confused compared to correctly rejected. Due to the generally low and inter-individually varying amount of *stream confusions*, this analysis may have lacked sufficient statistical power for the permutation test to reveal a significant cluster.

To further investigate whether the pupil-behavioral-correlation found earlier was merely driven by the increased response to *stream confusions* (which weaker performing subjects had more of), or also by a more general, trait-like correlation, we calculated a similar correlation, but this time based on responses to *hits* and *correct rejections* only (Fig. 3F). Convincingly, the correlation coefficient remained nearly identical (r = 0.74, CI_r_ = [0.453; 0.888], p = 1.25×10^-4^). This suggests that this correlation was not driven by the variability in the amount of *stream confusions* but by a general relationship between *pupil selectivity* and *behavioral selectivity*.

### Control analysis

#### Confusion of task instruction as potential cause for stream confusions

A control analysis investigated whether *stream confusions* were caused by mistakenly considering the irrelevant side as the relevant one, which should result in clustering of stream confusions and thus deviation from a random surrogate distribution (Fig. 4A). The random permutation did not result in significant deviations from the original distribution in any of the four bins counting the number of stream confusions (zero: p = 0.749, one: p = 0.595, two: p = 0.291, three: 0.607). This indicates that the main cause of stream confusion was momentary distraction rather than confusion of secondary task instructions.

**Figure 4:**
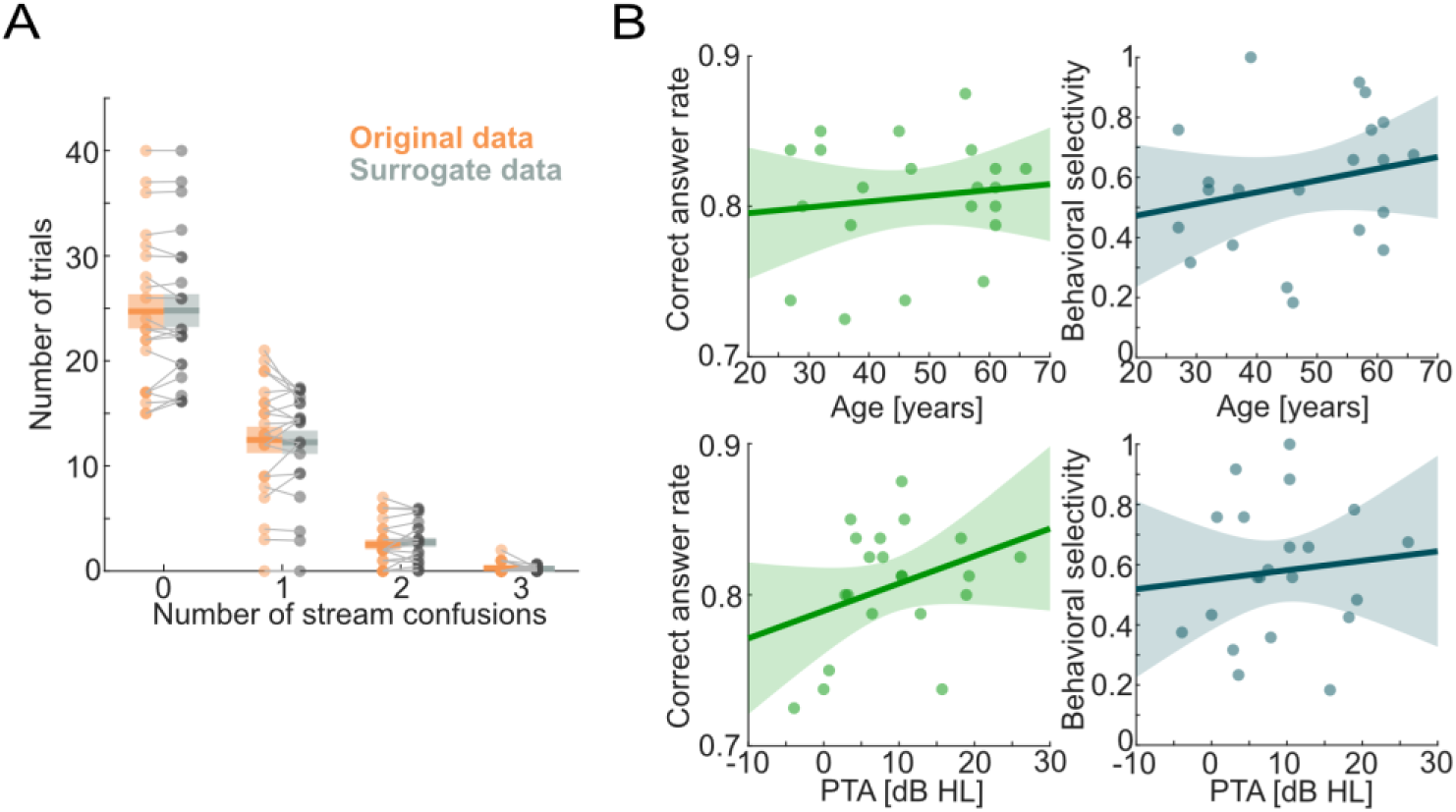
Control analysis: **A)** Control analysis: Histogram of number of stream confusions per trial for original data (orange) and surrogate data (grey) from a random permutation. **B)** Correlation of both age and PTA with both correct answer rate and behavioral selectivity.

**Figure 5:**
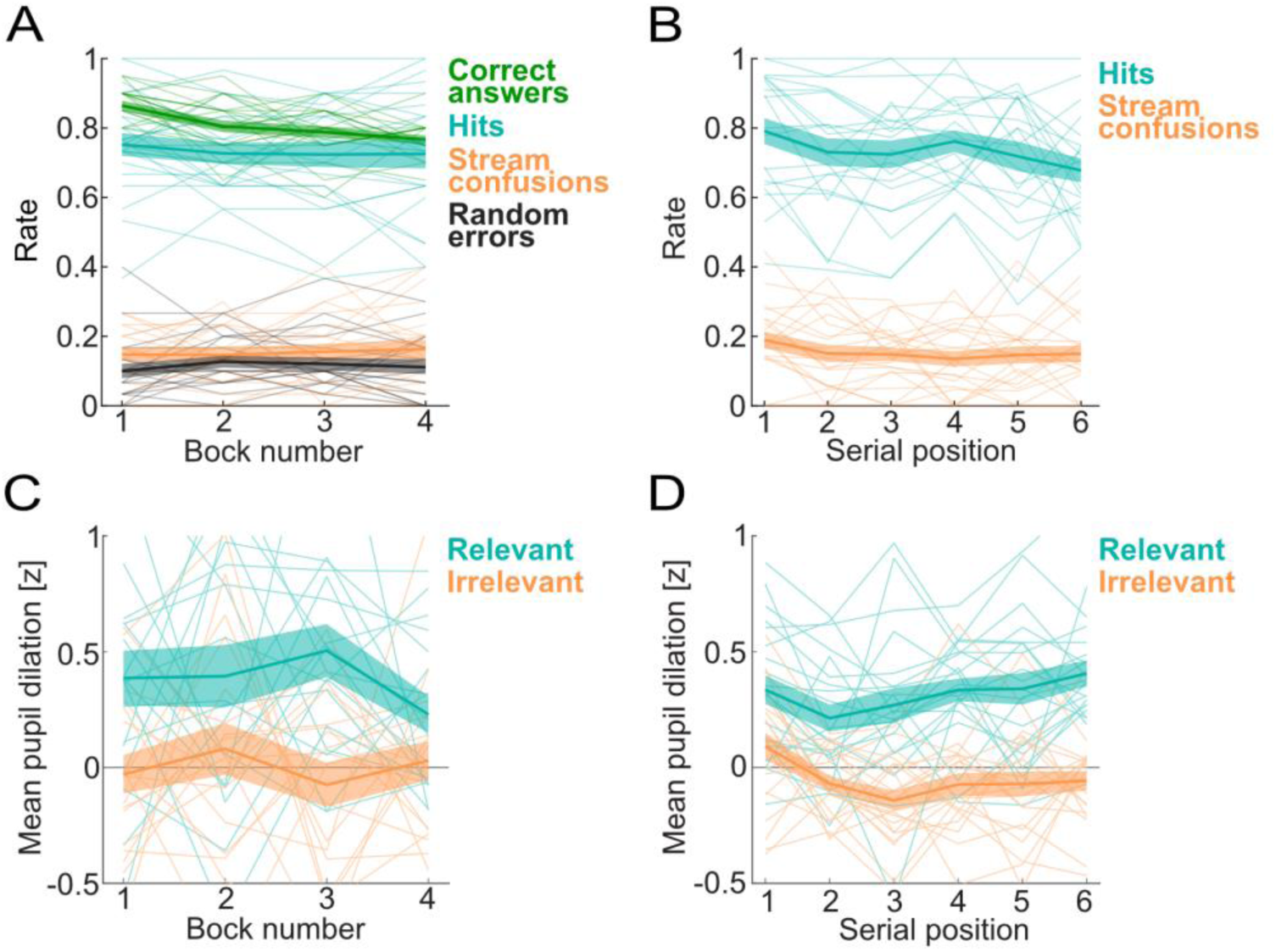
Serial order effects. Thick solid lines embedded in shaded areas depict the mean ±1 standard error. Thin lines depict individual means. **A)** Behavioral performance across the four experimental blocks (10 trials each). **B)** Proportion of hits and stream confusions per serial position within trial. **C)** Mean pupil dilation across the four experimental blocks. **D)** Mean pupil dilation across serial position within trial.

#### Correlation among age, hearing loss and behavior

To characterize the data on a descriptive level and to detect potential confounds, relationships between task performance, age, and PTA were analyzed and are depicted in Fig. 4B. However, neither correct answer rate nor behavioral selectivity correlated strongly with age (Correct answer rate: r = 0.127, CI_r_ = [-0.322; 0.53], p = 0.583; Behavioral selectivity: r = 0.232, CI_r_ = [-0.221; 0.604], p = 0.31) nor PTA (Correct answer rate: r = 0.335, CI_r_ = [-0.114; 0.66], p = 0.138; Behavioral selectivity: r = 0.105, CI_r_ = [-0.343; 0.513], p = 0.652). Since we controlled for individual variability in pupil size (see methods, above) and since behavioral selectivity did not show strong correlations with age or PTA, we did not include age or PTA as covariates in our pupil-behavioral analyses.

#### Time on task effects on behavior and pupil response

To investigate whether the effects may depend on time on task, both across the experiment (i.e., block number) and across a trial (i.e., serial position with trial), we investigated if slopes fitted to the individual data deviated from zero.

Across blocks, the negative slope of *correct answer rate* significantly differed from zero, indicating that participants got worse over the course of the experiment (mean slope = −3.1% per block, SD = 2.37%, t(20) = −5.99, p = 7.48×10^-6^, Cohens d = −1.31). There was no significant deviation from zero in the secondary task performance slopes, indicating that participants performance was rather constant (hits: p = 0.383; stream confusions: p = 0.407; random errors: p = 0.599).

MPD slopes across the blocks did not significantly differ from zero, indicating that *pupil selectivity* was rather constant (relevant: p = 0.401, irrelevant: p = 0.946).

Within trials, the negative slope of *hits* significantly differed from zero, indicating that earlier relevant background sounds are better memorized than later ones (mean = −1.61%/position, SD = 2.59%, t(20) = −2.856, p = 0.0098, Cohens d = −0.622). The negative slope of stream confusions just failed to reach statistical significance (mean = −0.64%/position, SD: 0.0165, t(20) = - 1.783, p = 0.0898).

Across serial position within trial, the positive slope in MPD of the response to relevant background sounds significantly differed from zero, indicating that later relevant background sounds led to stronger pupil responses than earlier ones (mean = 0.0228, SD = 0.0420, t(20) = 2.49, p = 0.0217, Cohens d = 0.543). The negative slope in MPD of the response to irrelevant background sounds was close to a significant difference from zero (mean: −0.093, SD = 0.0489, t(20) = −1.81, p = 0.0858, Cohens d = −0.395).

## Discussion

We asked whether pupil responses in an auditory attentional control task reveal relevance-related selectivity. Our findings indicate that such selectivity exists and predicts behavioral performance. The behavioral fate of task-irrelevant background sounds was partly indicated by pupil size, since stream confused irrelevant sounds led to greater pupil dilation than correctly rejected ones. Additionally, we found that both behavioral performance and pupil responses were relatively constant throughout the experimental session.

### Pupil responses to irrelevant background sounds indicate distraction

Correctly rejected irrelevant background sounds did not lead to clear pupil responses. This was surprising, given that earlier studies have shown that irrelevant sounds do indeed elicit pupil responses (Cronin et al., 2023; Fiedler et al., 2025; Hebisch et al., 2024; Marois & Vachon, 2018; Petersen et al., 2017; Tona et al., 2016). A likely reason for the absence of such a response is acoustic predictability of the background sounds: Names and numbers were all spoken by the same female voice. While uncertainty or novelty of sounds has been found to drive pupil responses (Friedman et al., 1973; Marois et al., 2020), highly predictable irrelevant stimuli are less distracting and lead to weaker pupil dilation (Vachon et al., 2012). Even though the exact timing of the name onset was unpredictable, spatial attentional filtering may have quickly adapted to the spatial task, such that most of the irrelevant background sounds could be easily inhibited. This raises an important question: to which extent is stimulus-driven attention actually automatic and how quickly can it be adapted (i.e., over the course of minutes) to the agenda of voluntary attention. Importantly, we found indications of a greater pupil dilation in response to stream confused irrelevant sounds. Stream confusions occurred over the course of the whole experiment, which means they could not be fully avoided by training of the voluntary attentional goals. In sum, our results confirm that pupil responses to irrelevant sounds indicate momentary distraction (Fiedler et al., 2025).

### Pupil responses to relevant sounds are a superposition of multiple responses

The observed pupil response to relevant background sounds is likely a complex of superimposed phasic and tonic components (Aston-Jones & Cohen, 2005). A phasic response to discrete items that require cognitive processing usually peaks at around 1.5 seconds before returning back to baseline (Hoeks & Levelt, 1993), which fits both the peaks following the name and the number in our data. However, a mere phasic response would also return to baseline after around 3 seconds, while we observed a longer-lasting sustained dilation, which we hence consider rather a tonic component.

First, pupil size increased in response to the name. While the name itself was task-irrelevant, it served as a cue for the upcoming number. The associated pupil dilation may reflect the effort of shifting attention towards the respective side (McCloy et al., 2017, 2018) as well as anticipating the effort need to process the upcoming number (Alfandari et al., 2023; Irons et al., 2017; Unsworth & Miller, 2023).

Second, the pupil dilation in response to the number may reflect both the actual listening effort associated with comprehension and encoding into memory. Since participants were asked to memorize all stimuli from the relevant side, we cannot disentangle listening effort from memory encoding. A holistic view may even count memory retrieval under listening effort (Pichora-Fuller et al., 2016), since it is an essential element of the listening task. Future studies may be able to disentangle comprehension from memory encoding by introducing an alternative secondary task instruction that depends on comprehension: For example, participants could be instructed to memorize odd (or even) numbers, such that comprehension is needed before a background sound can be identified as relevant or irrelevant.

Third, we observed a rather tonic plateau of the pupil size staying above baseline. This plateau could be explained by either the general memory load or the effort exerted for active memory maintenance. A memory-load related pupil dilation has been found earlier (Goldinger & Papesh, 2012; Kahneman & Beatty, 1966), also in speech-in-noise tasks (Bönitz et al., 2021; Micula et al., 2022). If general memory load would lead to such a plateau, falsely memorized irrelevant numbers (i.e., stream confusions) should also elicit a similar response, but we did not observe that. This suggests that effort exerted for active memory maintenance is more likely to have caused this plateau. Participants may have rehearsed the secondary targets occasionally even though they were asked to fully switch attention back to the primary target after memory encoding of a secondary target. Interestingly, this memory maintenance did not seem to be a warranty for a later hit as indicated by the absence of a difference between pupil responses to hits and misses. This may speak against the memory maintenance hypothesis, but we think it rather indicates that the main obstacle in this task was not the allocation of sufficient attentional resources to relevant stimuli but rather the inhibition of irrelevant stimuli.

### Pupil response selectivity reflects auditory attentional control

We found task-relevance related selectivity in the pupil response and that the strength of the selectivity correlates with behavioral performance. Attention-related selectivity in pupil responses has been observed before, mainly in the visual domain (Binda et al., 2013, 2014; Mathôt et al., 2013, 2014; Tkacz-Domb & Yeshurun, 2018) but also in the auditory domain (Fiedler et al., 2025; Liao et al., 2016; Pielage et al., 2021). The tight link to behavioral performance under an auditory attentional control paradigm with relatively high ecological validity (e.g., in comparison to an oddball paradigm) is novel to our knowledge. Since we found that this selectivity is even present when we controlled for the behavioral fate of the background sounds, we conclude that the pupil response selectivity reflects not only momentary, state-like distraction, but also trait-like auditory attentional control abilities. Note that even though we found a rather strong correlation coefficient, the relatively small sample size does not allow us to draw strong conclusions about the true correlation (Schönbrodt & Perugini, 2013). However, with the lower bound of the confidence interval at .45 we can at least conclude that the true correlation probably has a considerable strength.

One limitation of our study is that we have not conducted any cognitive tests, which could indicate to which extent general cognitive abilities explain auditory attentional control. Attentional control abilities are usually assessed in the visual domain, for example with the Stroop task (Stroop, 1935) or the flanker task (Eriksen & Eriksen, 1974), also using pupillometry (for review: Robison, 2024). Furthermore, working memory capacity likely plays a strong role when it comes to memorizing two-digit numbers.

### What does pupillometry reveal beyond behavioral performance?

From a physiological point of view, the findings presented here provide insight into how the pupil is linked to attentional processes. From a clinical perspective, however, one might question whether pupillometry is merely a reflection of behavior or if it can offer additional insights into the individual challenges related to attentional control. It has been shown earlier that pupil size reveals challenges that are not reflected in behavior, but rather in additional effort exerted in compensatory mechanisms. For example, under decreasing SNR in a speech-in-noise test, the pupil response starts to increase before behavior is affected (Ohlenforst et al., 2018). Retrospective mental repair that maintains correct behavioral responses was shown to increase pupil responses (M. B. Winn & Teece, 2021). This suggests that some effects only manifest in behavior if the experimental manipulation of task load is driven into unrealistically difficult conditions, such as negative SNRs in speech-in-noise tests (Smeds et al., 2015), whereas pupillometry is able to reveal these challenges under more ecologically valid conditions.

In the present study, behavioral performance provided insights into the inter-individual variability of attentional control abilities. However, only through pupil size could we infer that the predominant challenge was inhibition of irrelevant background sounds (see above). Furthermore, pupillometry may allow to study subtle differences in how much attention is captured in a stimulus-driven fashion due to the acoustic properties of a sound, which may not have an immediate impact on behavior (Fiedler et al., 2025). In sum, pupil size under attentionally challenging conditions may reveal intra- and interindividual differences that would only manifest in behavior if driven to the extreme.

As argued in the introduction, one potential field of clinical application is hearing aid fitting: Besides the frequency-specific amplification to compensate for elevated hearing thresholds, a hearing care professional needs to adjust the strength of noise reduction and directionality to ideally match the hearing-aid users’ abilities and preferences. While recently introduced diagnostic measures (Zaar et al., 2024) certainly improved the individualization of noise reduction and directionality, they still only target speech-in-noise performance and do not take the individual attentional control abilities into account. Our paradigm presented here, including pupillometry, can help evaluate to what extent the manipulation of noise reduction and directionality interacts with individual attentional control abilities and whether this interaction can be predicted from domain-general tests of attentional control. Furthermore, data gained in such a paradigm can be used to analyze which psychoacoustic sound features are beneficial for attentional control: There may be certain sound features that mark relevance (e.g., the spectro-temporal modulation of an ambulance sound), and others are typically irrelevant (e.g., the constant noise of an air condition). Based on such an analysis, hearing aids could be fine-tuned to amplify typical relevant features and to attenuate irrelevant ones.

## Conclusion

In an auditory attentional control task, we found that the pupil response shows task-relevance related selectivity, which is closely linked to behavioral performance. We interpret the pupil responses as the amount of attentional resources deployed to shift, comprehend, and memorize background sounds, which could be predominantly observed in response to relevant, but also in response to stream-confused irrelevant background sounds. We conclude that the pupil response provides insights into the individual challenges in attentional control, which can be of clinical relevance, for example in the development of diagnostic tools for personalization of noise management in hearing aids.

## Data availability statement

The data and material is available upon request based on a data sharing agreement to comply with the participants’ consent.

## Author Contribution

L.F. designed the study, implemented experimental procedures, collected data, analyzed the data, and wrote the manuscript. I.J. and D.W. designed the study and revised the manuscript.

## Acknowledgements

Garghi Seenevas supported the implementation of experimental procedures, data collection, and analysis. Torben Christiansen (EPOS Group A/S) contributed to the study design with discussion of potential application in noise-cancelling headsets.

